# Murine modeling of IDH-mutant 1p/19q-codeleted oligodendroglioma reveals genotype specific phenotypes

**DOI:** 10.64898/2026.05.14.725183

**Authors:** Ipsita G. Kundu, Nicolas Toro, Aaron Y. Mochizuki, Jaldeep Langhnoja, Rithvik V. Ayyagari, James C. Cronk, Sarah M. Reel, Phonepasong Arounleut, Luis Tron Esqueda, Christine E. Fuller, Pavithra Viswanath, Amy B. Heimberger, Craig M. Horbinski, Timothy N. Phoenix

## Abstract

Oligodendroglioma is a primary central nervous system tumor classified by the presence of isocitrate dehydrogenase (IDH) mutations and codeletion of 1p/19q. Here we describe the generation of an IDH-mutant 1p/19q-codeleted oligodendroglioma mouse model using in utero electroporation. We identified IDH1^R132H^, PIK3CA^E545K^, *Cic*^KO^, *Fubp1*^KO^ and *Cdkn2a*^KO^ as the optimal combination (termed Oligo^Cdkn2a^) to drive fully penetrant tumors that histologically resemble human grade II/III IDH-mutant, 1p/19q-codeleted oligodendroglioma. Replacing *Cdkn2a* with *Trp53* loss in this mouse model shifted tumor histology towards high grade astrocytoma. Oligo^Cdkn2a^ tumors displayed metabolic and transcriptional changes associated with *IDH* and *CIC* mutations, and single cell sequencing identified a bias towards oligodendrocyte differentiation compared to an IDH wild-type glioblastoma mouse model. Oligo^Cdkn2a^ tumors represent the first mouse model system to recapitulate the genetic, histological and transcriptional features of human IDH-mutant 1p/19q-codeleted oligodendrogliomas, offering a platform to further dissect tumor biology and test new therapeutic strategies.

## Introduction

IDH-mutant 1p/19q-codeleted oligodendroglioma (termed IDH-O) is a subtype of adult-type diffuse glioma^1^. IDH-O tumors present as grade 2/3, with peak incidence occurring between 35-56 years old^2,3^. While IDH-O has a better prognosis compared to other types of adult-type diffuse glioma^4,5^, the current standard of care treatments including surgery, radiotherapy and alkylating agents significantly impact patient quality of life^6,7^. Vorasidenib has recently been approved by the FDA for treating grade 2 IDH-mutant gliomas based on trials that demonstrated significant improvement in progression free survival and delays the need for the next intervention^8–10^. However, patients with grade 3 or 4 IDH-mutant and/or with gadolinium enhancement do not respond as favorably. The ability to better understand IDH-O and develop new therapies has been hindered by a lack of available model systems^5^.

IDH-mutations, which serve as a defining molecular hallmark for both IDH-O and IDH-mutant astrocytoma (termed IDH-A), lead to increased production of D-2-hydroxyglutarate (D-2HG) which is a competitive inhibitor of alpha-ketoglutarate (α-KG) dependent dioxygenases, including DNA and histone demethylases^11–15^. Accumulation of D-2HG drives widespread epigenetic remodeling characterized by global DNA hypermethylation and subsequent transcriptional reprogramming including the aberrant activation of *PDGFRA* and silencing of *CDKN2A*^16,17^. In parallel, the consumption of α-KG and NADPH by mutant IDH enzyme promotes metabolic reprogramming, and multiple studies have uncovered selective vulnerabilities of IDH-mutant gliomas to pathways including oxidative phosphorylation (OXPHOS)^18^, glutaminase^19^, branched-chain amino acid transaminase^20^, and nicotinamide phosphoribosyltransferase (NAMPT)^21^.

Now defined by the combination of IDH mutation and 1p/19q codeletion, IDH-O exhibits several distinguishing features that separate them from IDH-A and IDH wild-type glioblastoma (GBM). IDH-O are histologically characterized by relatively uniform tumor cells with round nuclei and fast development of hydropic cytoplasmic changes after surgical removal, imparting a classic “fried egg” appearance^1,22^. Beyond 1p/19q codeletion, IDH-O are enriched for loss-of-function alterations in *CIC* (located on Chr19q), *FUBP1* (located on Chr1p) and *NOTCH1;* whereas *TP53* mutations, which are characteristic of IDH-A and GBM, are typically absent^23–28^.

Comparative transcriptomic analyses of IDH-A and IDH-O demonstrate that their biological divergence is driven by these distinct genetic alterations. Single cell RNA-sequencing of IDH-mutant diffuse gliomas has revealed a shared glial cell lineage program, with tumor-intrinsic transcriptional differences primary driven by genotype-specific events (e.g., *CIC* vs. *TP53* alterations)^29^. Notably, differences found in bulk expression profiles are derived from the tumor microenvironment (TME). For example, IDH-A is associated with increased immune cell infiltration and activation relative to IDH-O^30,31^. Thus, the distinctive genetic landscape of IDH-mutant diffuse glioma shapes not only their tumor cell-intrinsic transcriptional programs, but also the other populations within the TME.

Animal models that faithfully recapitulate the genetic, pathological, and transcriptional heterogeneity of adult-type diffuse gliomas are essential tools for biological discovery and preclinical therapeutic evaluation. Multiple genetically engineered mouse models (GEMMs) for GBM have been established, yielding insight into cellular origins, mechanisms of oncogenic drivers, TME interactions and responses to candidate therapies^31–33^. Mouse models of IDH-A driven by combinations of *Trp53* and *Atrx* loss^34^ and/or oncogenic *RasV12* expression have identified IDH-mutant specific vulnerabilities, alterations in TME interactions^35^, and mechanistic dissection of glioma-associated risk variants at 8q24.21 that confer an approximately sixfold increased risk of developing IDH-mutant glioma^36^. Despite these advances, there are no GEMMs that faithfully represent IDH-mutant 1p/19q-codeleted oligodendroglioma, limiting mechanistic interrogation and preclinical testing in this genetically and biologically distinct subtype.

Here, we detail the generation of IDH-O mouse models. Using in utero electroporation (IUE) to introduce complementary gain- and loss-of-function mutations into the developing embryonic cortex, we define genetic combinations that drive fully penetrant tumor formation. IDH-O mouse models with *Cdkn2a* loss display the longest latency and best recapitulate the key features of human IDH-O tumors, whereas those with *Trp53* loss exhibit astrocytoma histology and more aggressive behavior. We further demonstrate metabolic reprogramming in IDH-O mouse models relative to an IDH wild-type GBM mouse model and identify p53 status as a modulator of TME interactions. Together, these models provide a versatile platform to dissect how genetic context impacts tumor pathogenesis and will enable preclinical evaluation of targeted and immune-modulating therapeutic strategies for high-grade IDH-O.

## Results

### Generation of IDH-mutant 1p/19q-codeleted oligodendroglioma mouse models

To model IDH-mutant 1p/19q-codeleted oligodendroglioma, we prioritized the most frequent genetic alterations identified in patient samples^23–25^ (**Supplementary Figure 1a**) that could be recapitulated in the developing mouse brain using in utero electroporation (IUE). These included the expression of *IDH1*^R132H^ and *PIK3CA*^E545K^ along with loss of the respective 19q and 1p targets *Cic* and *Fubp1* (termed Oligo). PiggyBac plasmid and CRISPR-Cas9 components for this Oligo combination were electroporated at embryonic day 14.5 (E14.5) and viable electroporated pups were monitored for up to 200 days post-birth (**Figure 1a**). Oligo mice did not develop neurological symptoms related to tumor burden, and examination of brains harvested at early, mid and late time points (30, 90 and ∼200 days post-birth respectively) showed no overt signs of tumor formation (**Figure 1b**).

**Figure 1.**
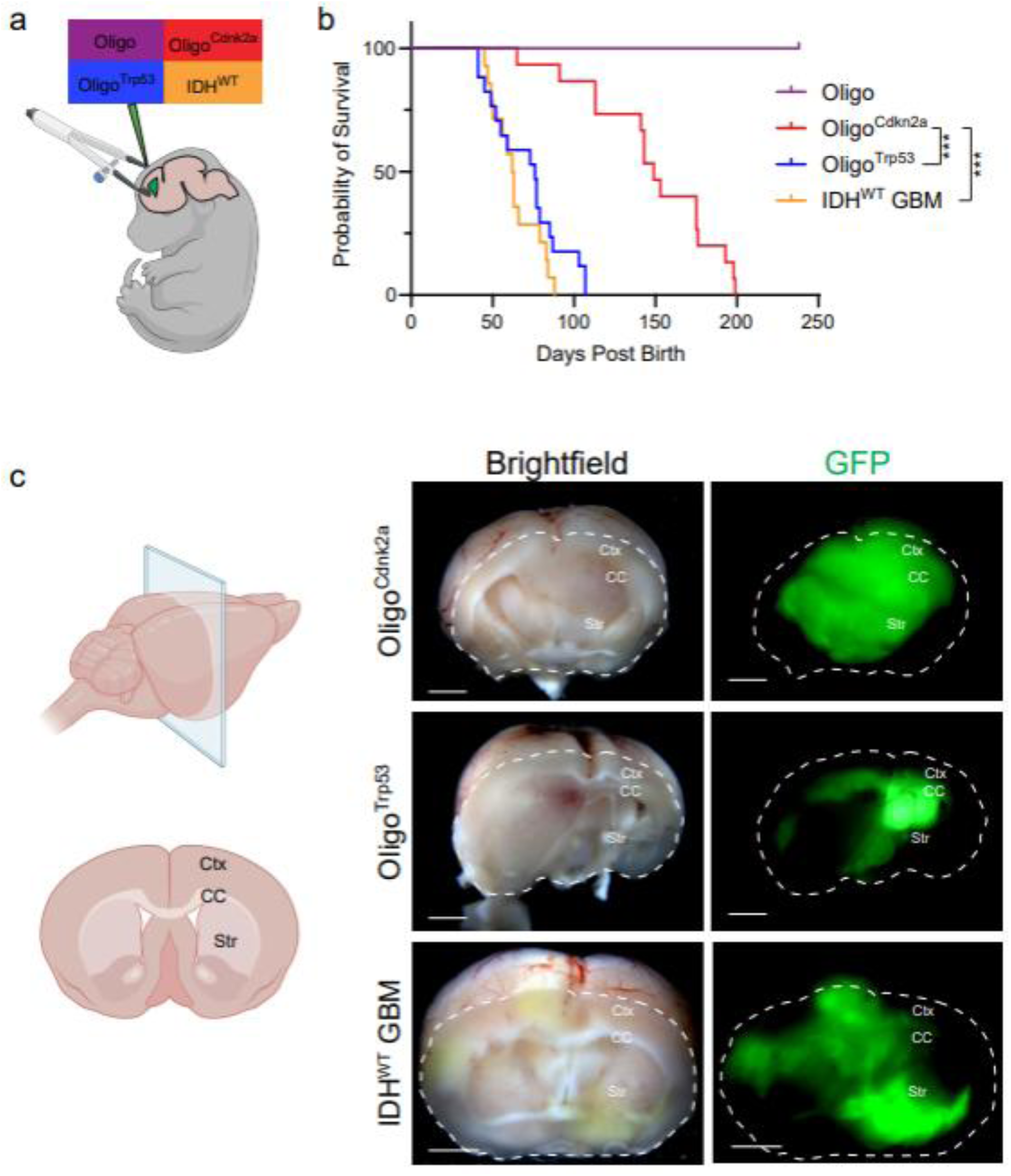
Generation of IDH-mutant 1p/19q-codeleted oligodendroglioma mouse models. **a,** Schematic diagram of IUE conditions tested. **b,** Kaplan Meier survival curves for indicated IUE conditions. Oligo (n=16), Oligo^Cdkn2a^ (n=15) median survival 149 days, Oligo^Trp53^ (n=17) median survival = 76 days, IDH^WT^ GBM (n=14) median survival 62.5 days. ***p<0.0001, log rank mantel cox test. **c,** Schematic diagram of mouse brain orientation and representative brightfield and GFP images demonstrating tumor location in coronal brain section from indicated IUE conditions. Scale bar = 200µm. Ctx: cortex; CC: corpus callosum; Str: striatum.

To determine if the concomitant loss of a tumor suppressor is required for tumorigenesis, we created two additional models by CRISPR-Cas9 targeting of *Cdkn2a* (termed Oligo^Cdkn2a^) or *Trp53* (termed Oligo^Trp53^) in the original Oligo IUE combination. IUE of Oligo^Trp53^ drove the formation of fully penetrant tumors with a median survival of 76 days post-birth (n=17) (**Figure 1b**). GFP-positive tumor cells could be found in the corpus callosum region and frequently exhibited hemorrhagic areas (**Figure 1c** and **Supplementary Figure 1c**). IUE of Oligo^Cdkn2a^ also induced fully penetrant tumors, but with significantly longer latency at a median survival of 149 days post-birth (n=15) (**Figure 1b**). Oligo^Cdkn2a^ tumors were similarly localized to the corpus callosum region but did not commonly display hemorrhagic regions like those observed in Oligo^Trp53^ (**Figure 1c** and **Supplementary Figure 1b**).

For comparison, we also generated an established murine model of IDH-wildtype glioblastoma through IUE mediated CRISPR-Cas9 disruption of *Nf1*, *Pten,* and *Trp53* (termed IDH^WT^ GBM)^37,38^. IDH^WT^ GBM IUE produced tumors with a median survival of 63 days post-birth (n=14) (**Figure 1b**). IDH^WT^ GBM tumors were highly invasive with GFP-positive tumor cells present throughout the cerebrum and even found in distant brain structures (e.g., brainstem) (**Figure 1c** and **Supplementary Figure 1d**). Together, these data demonstrate that the introduction of recurrent mutations found in IDH-O patients, when combined with the additional loss of a tumor suppressor, is sufficient to drive the formation of fully penetrant brain tumors *in vivo*.

### Oligo^Cdkn2a^ mouse models recapitulate histopathological features of human IDH-mutant 1p/19q-codeleted oligodendroglioma

Neuropathological assessment of murine tumors revealed distinct genotype-specific growth patterns and cytologic phenotypes. Oligo^Cdkn2a^ tumors extended into the overlying cortex but preferentially tracked along the corpus callosum (**Figure 2a**). At higher magnification, tumors resembled WHO grade 2–3 oligodendroglioma with scattered mitoses and rounded nuclei with relatively open chromatin. The classic “fried egg” perinuclear halo, a hallmark feature of human oligodendrogliomas, was also prominent in Oligo^Cdkn2a^ samples (**Figure 2b**). By contrast, Oligo^Trp53^ tumors were more aggressive and highly angiogenic, expanding to occupy most of the ipsilateral hemisphere without substantial contralateral spread (**Figure 2a**). Notably, Oligo^Trp53^ tumors displayed angulated nuclei with denser chromatin and overtly astrocytic cytology (**Figure 2b**). This suggests that *Trp53* loss drives a shift in nuclear morphology. IDH^WT^ GBM produced the most diffusely infiltrative phenotype, with tumor cells infiltrating throughout the ipsilateral hemisphere, crossing the midline into the contralateral hemisphere, and disseminating to multiple distant regions (**Figure 2a**). Further analysis of IDH^WT^ GBM showed numerous mitoses, regions of necrosis, and markedly enlarged epithelioid-like tumor cells with clear borders (**Figure 2b**), resembling epithelioid variants observed in GBM patient samples.

**Figure 2.**
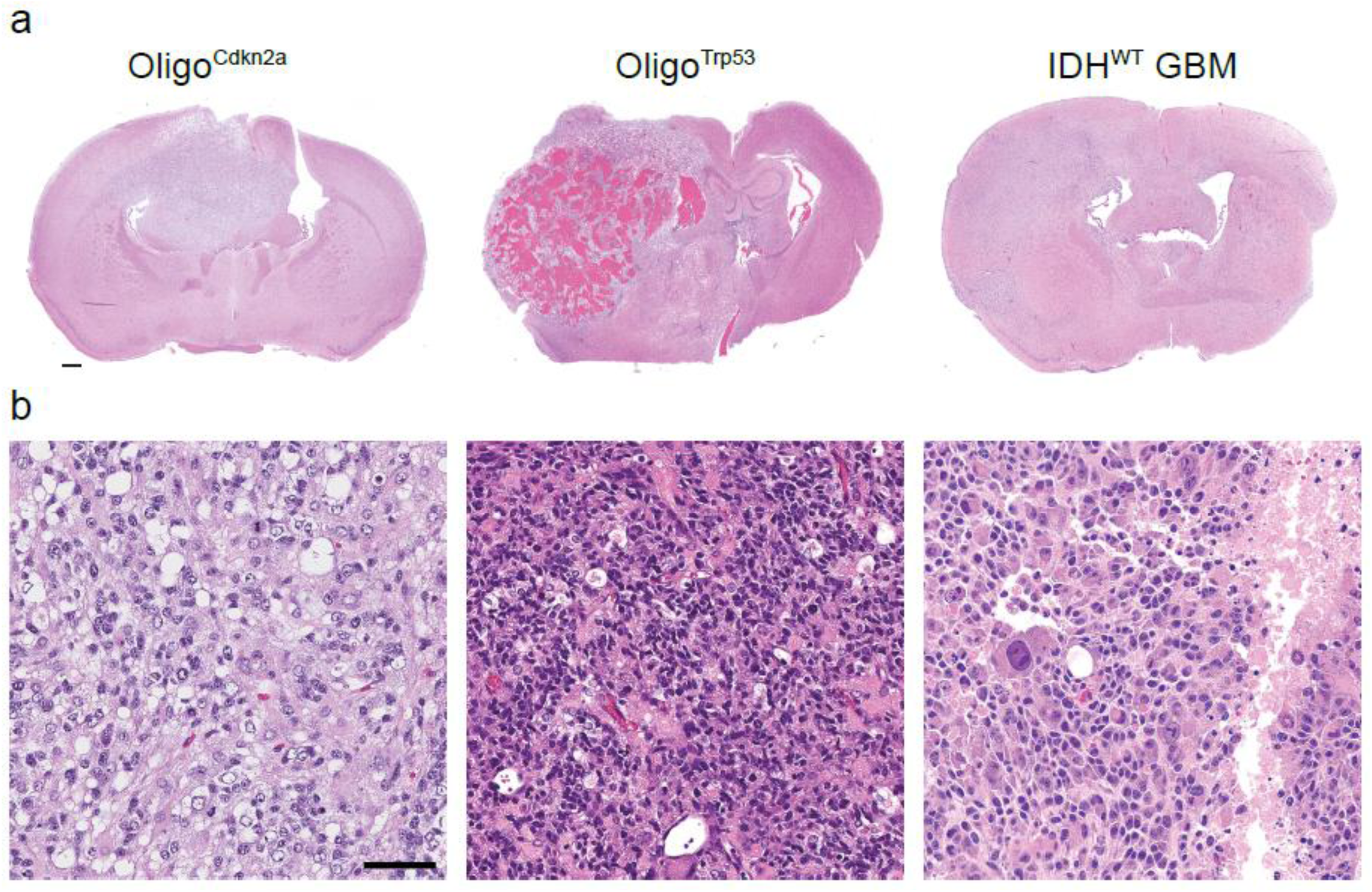
Oligo^Cdkn2a^ mouse models recapitulate the histopathological features of human IDH-mutant 1p/19q-codeleted oligodendroglioma. **a,** Low-magnification overview of coronal H&E sections for each condition. All three IUE conditions showed diffusely infiltrative growth patterns (upper), with the Oligo^Trp53^ containing abundant neovascularization and mass effect (upper middle). **b,** At higher magnification, the Oligo^Cdkn2a^ tumors showed mostly rounded nuclei and hydropic change (lower left), while the Oligo^Trp53^ cells were more angulated with darker clumped chromatin (lower middle). IDH^WT^ GBM cells showed the most anaplasia, as well as necrosis (lower right). Scale bars: 2mm for low magnification images in (a) and 60µm for higher magnification images in (b).

Additional immunofluorescent staining across IUE tumor conditions showed co-labeling of GFP-positive tumor cells with the glioma-associated marker Olig2 (**Supplementary Figure 2a-c**). Ki67 immunoreactivity confirmed proliferative tumor cells across conditions, and staining for glial fibrillary acidic protein (GFAP) highlighted both astrocyte-like tumor cells and reactive astrocytes within tumors (**Supplementary Figure 2a-c**).

### Validation of gain- and loss-of-function mutations in Oligo^Cdkn2a^, Oligo^Trp53^, and IDH^WT^ GBM mouse models

Immunohistochemistry (IHC) for GFP across conditions mirrored tumor location and invasion patterns found at the time of brain collection and by neuropathological review (**Figure 3a**). IHC with an IDH1 R132H specific antibody showed overlapping staining patterns with GFP in Oligo^Cdkn2a^ and Oligo^Trp53^ conditions, while IDH^WT^ GBM samples were negative (**Figure 3a**). Using Proton magnetic resonance spectroscopy (1H-MRS)-based metabolite profiling, we readily detected the oncometabolite 2-HG in Oligo^Cdkn2a^ and Oligo^Trp53^ tumor cells, whereas 2-HG was undetectable in IDH^WT^ GBM cells, consistent with IDH-mutant restricted production (**Figure 3b**). Both Oligo^Cdkn2a^ and Oligo^Trp53^ tumor cells were responsive to vorasidenib, which reduced 2-HG levels to baseline (**Figure 3b**). 2-HG is known to inhibit pyruvate dehydrogenase activity, thereby reducing glucose flux to glutamate and its steady-state abundance^39^. Accordingly, treatment with vorasidenib normalized glutamate levels in Oligo^Cdkn2a^ and Oligo^Trp53^ tumor cells further verifying metabolic rewiring in these IDH-mutant models (**Supplementary Figure 3a-c**).

**Figure 3.**
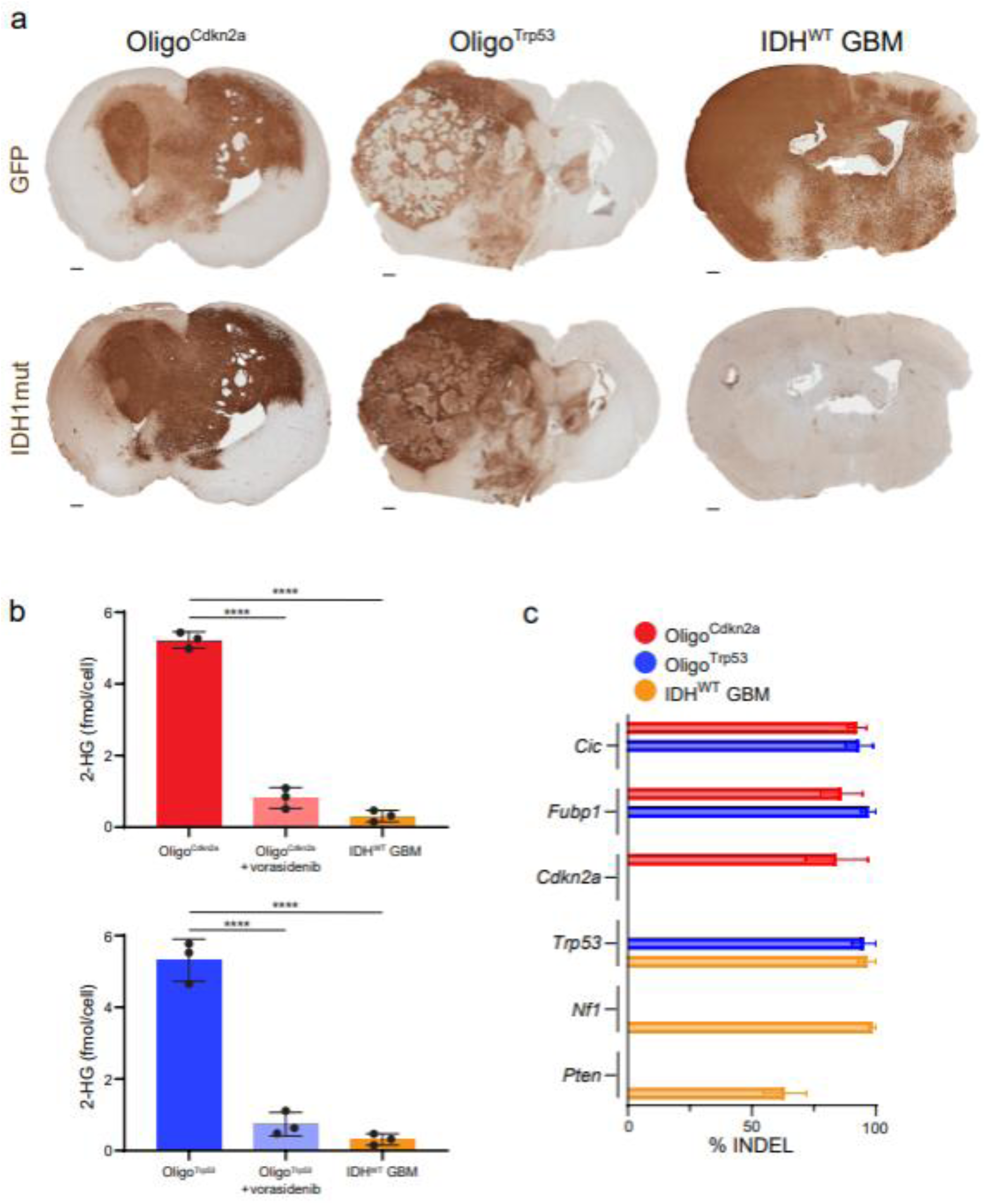
Validation of gain- and loss-of-function mutations in Oligo^Cdkn2a^, Oligo^Trp53^, and IDH^WT^ GBM mouse models. **a,** Low magnification overview of tumor coronal sections stained by IHC for GFP and IDH1-mutation. Scale bar: 500µm.**b,** 2-HG concentrations in Oligo^Cdkn2a^, Oligo^Trp53^, and IDH^WT^ GBM tumor cell conditions. One-way ANOVA with Tukey’s multiple comparisons test. ****p<0.0001. Values indicate mean ± SD; Oligo^Cdkn2a^ n=3, Oligo^Trp53^ n=3, IDH^WT^ GBM n=3. **c,** INDEL frequency for denoted gene target *Cic*, *Fubp1*, *Cdkn2a*, *Trp53*, *Nf1*, and *Pten* in tumor cells from each condition. Values indicate mean ± SEM; Oligo^Cdkn2a^ n=6; Oligo^Trp53^ n=5; IDH^WT^ GBM n=6.

Targeted sequencing of tumor cell DNA demonstrated efficient on-target editing of CRISPR–Cas9 loci in each IUE condition. *Cdkn2a*, *Cic* and *Fubp1* all displayed highly efficient editing in Oligo^Cdkn2a^ tumor cells, whereas Oligo^Trp53^ cells showed editing restricted to *Trp53*, *Cic* and *Fubp1* (**Figure 3c**). *Nf1*, *Pten,* and *Trp53* displayed high INDEL rates in IDH^WT^ GBM cells and were negative for editing in oligo-specific targets (**Figure 3c**). Collectively, the verified expression and knock-out of specific genetic combinations across IUE conditions demonstrates the ability to precisely model both gain- and loss-of-function alterations that drive glioma subtype development.

### IDH- and Trp53-mutation status impact the transcriptional profile of tumor models

Next, we performed whole-transcriptome sequencing (RNA-seq) on samples isolated from primary mouse tumors to investigate differences between genomic backgrounds. Principal component analysis (PCA) and Pearson’s correlation plots showed each condition segregated into independent clusters (**Supplementary Figure 4a, b**). K-means clustering of the top 2,000 differentially expressed genes identified three main clusters among tumor conditions (**Figure 4a**). Genes enriched in cluster 1 were upregulated in both *Trp53*-mutant conditions (IDH^WT^ GBM and Oligo^Trp53^) and were associated with gene sets related to immune system pathways. Cluster 2 genes were unique to IDH^WT^ GBM and enriched in neuronal pathways, whereas cluster 3 genes were predominantly upregulated in Oligo^Cdkn2a^ tumors and enriched in gene sets associated with extracellular matrix (ECM), focal adhesion, and PI3K-Akt signaling pathways (**Figure 4b**). Direct comparison of differentially expressed genes (FC >2; adj-p value < 0.05) between conditions using over-representation analysis further confirmed the specific enrichment of gene set pathways identified by K-means clustering (**Supplementary Figure 4c-e**).

**Figure 4.**
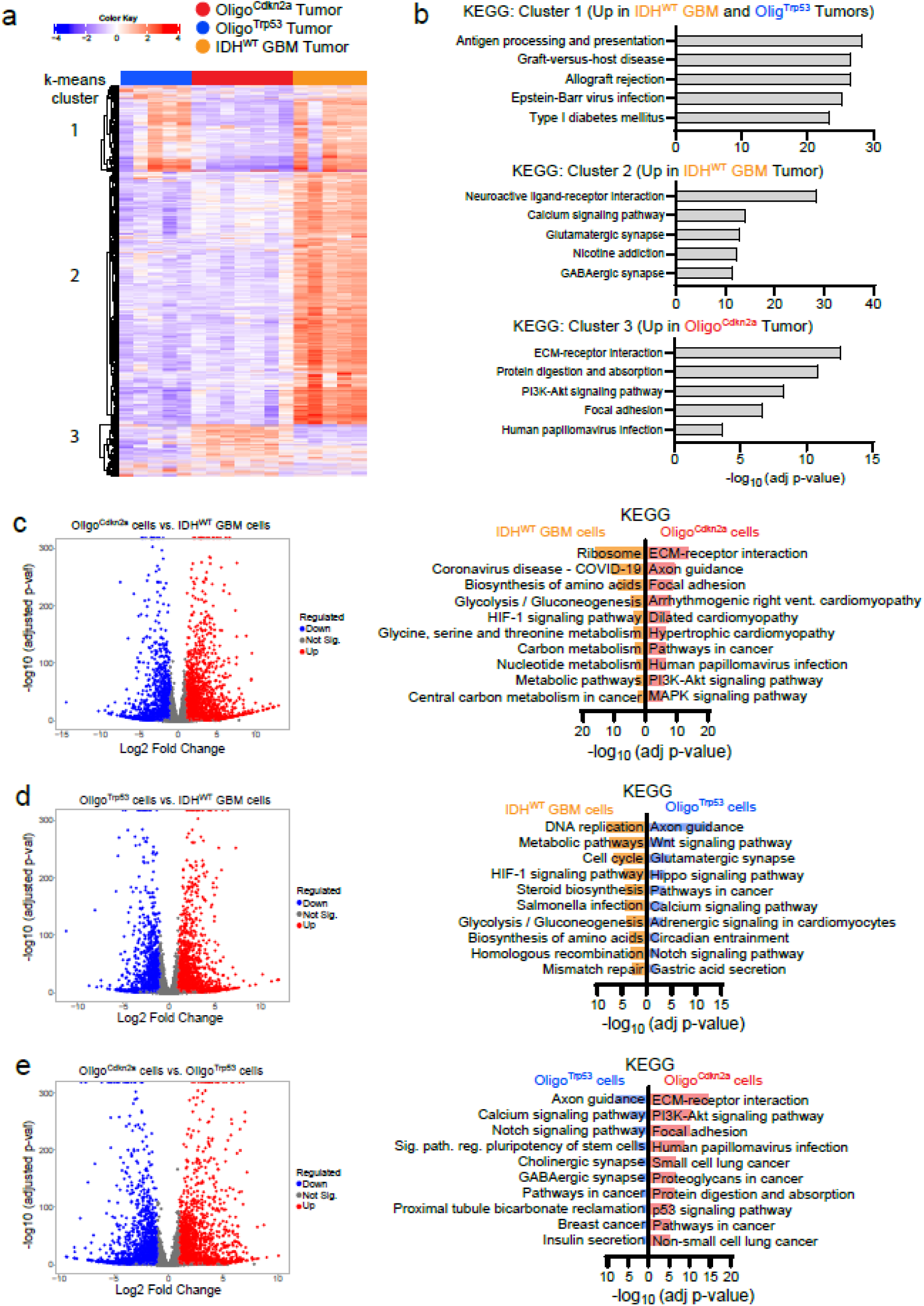
IDH- and Trp53-mutation status impact the transcriptional profile of tumor models. **a,** Heatmap of tumor conditions organized by K-means clustering. **b,** ORA of KEGG gene sets in each K-means cluster. The top five most significant gene sets are plotted by −log_10_(adj-pvalue). **c,** Volcano plot displaying significantly up-and down-regulated genes (FC ≥ 2 or ≤ −2 and adj p-val < 0.05) between Oligo^Cdkn2a^ vs IDH^WT^ GBM cells, and ORA of KEGG gene sets in each condition. The top five most significant gene sets are plotted by −log_10_(adj-pvalue). **d,** Volcano plot displaying significantly up- and down-regulated genes between Oligo^Trp53^ vs IDH^WT^ GBM cells, and ORA of KEGG gene sets in each condition. The top five most significant gene sets are plotted by −log_10_(adj-pvalue). **e,** Volcano plot displaying significantly up- and down-regulated genes between Oligo^Cdkn2a^ vs Oligo^Trp53^ cells GBM cells, and ORA of KEGG gene sets in each condition. The top five most significant gene sets are plotted by −log_10_(adj-pvalue).

We next conducted additional RNA-sequencing on cell lines derived from the mouse tumors to examine tumor cell specific transcriptional programs. IDH-mutations are known to induce metabolic reprogramming and are associated with reduced glycolysis relative to IDH^WT^ GBM^40–42^. When comparing IDH^WT^ GBM cells with Oligo^Cdkn2a^ or Oligo^Trp53^ cells, cellular metabolism, glycolysis/gluconeogenesis, and biosynthesis of amino acids were among the top KEGG pathways overrepresented in IDH^WT^ GBM cells, highlighting the impact of mutant IDH1 expression on tumor cell metabolic programs (**Figure 4c, d**). Axon guidance pathway genes were enriched in both Oligo^Cdkn2a^ and Oligo^Trp53^ cells when compared to IDH^WT^ GBM cells (**Figure 4c, d**). Additional pathways enriched in Oligo^Trp53^ cells included several developmental signaling pathways (e.g., Wnt and Notch pathways), while Oligo^Cdkn2a^ cells showed increased activation of ECM, focal adhesion, PI3K-Akt, and p53 signaling pathways (**Figure 4c-e**). Together, these data validate the functional consequence of mutant IDH1 expression on the metabolic programs of IUE brain tumor models, reveal activation of neuronal, ECM/adhesion, and PI3K/Akt pathway in Oligo^Cdkn2a^ and Oligo^Trp53^ models, and identify immune system pathway activation in *Trp53*-mutant tumor conditions.

### Single cell profiling identifies tumor cell-state and transcriptional heterogeneity driven by tumor genetics

To determine more granular differences in the cellular composition and associated cell-state programs of tumors, single cell transcriptional profiling across Oligo^Cdkn2a^, Oligo^Trp53^ and IDH^WT^ GBM models was performed. A total of 14 tumors samples including six Oligo^Cdkn2a^, four Oligo^Trp53^ and four IDH^WT^ GBMs were collected at endpoint and processed using the 10x GEM-X Flex platform. After filtering for quality control metrics, a total of 138,426 cells were used for downstream analyses. Cell-type identity of clusters were assigned by established gene expression markers, identifying tumor cell and non-tumor cell compartments (astrocyte, oligodendrocyte, endothelial, mural, myeloid, lymphoid, and ependymal/choroid plexus) (**Figure 5a, Supplementary Figure 5a**). Re-clustering of tumor cells alone identified cell-state programs previously described in human gliomas. These included clusters expressing genes associated with the cell cycle (Cycling-G2/M, Cycling-S1 and Cycling-S2), neural progenitor cells (NPC-like), oligodendrocyte progenitor cells (OPC-like-1 and OPC-like-2), oligodendrocyte differentiation and maturation (OL-like-early, OL-like-mid and OL-like-late), astrocytes (AC-like) and mesenchymal cells (Mes-like) (**Figure 5b** and **Supplementary Figure 5c, d**). The OL-like-mid and OL-like-late clusters displayed the most prominent differences between conditions, with both Oligo^Cdkn2a^ and Oligo^Trp53^ tumors containing increased proportions of these cells compared to IDH^WT^ GBM (**Figure 5c**). Other significant shifts in tumor cell composition included increased Cycling-G2/M and Cycling-S1 cells in Oligo^Trp53^ tumors, increased OPC-like-1 cells in Oligo^Cdkn2a^ tumors, and a trend for increased OPC-like-2 cells in Oligo^Trp53^ and IDH^WT^ GBM tumors (**Figure 5c**). These findings indicate that Oligo^Cdkn2a^ and Oligo^Trp53^ genotypes are uniquely permissive for tumor cells to differentiate along the oligodendrocyte trajectory, while *Trp53* loss supports the maintenance of cycling progenitor programs.

**Figure 5.**
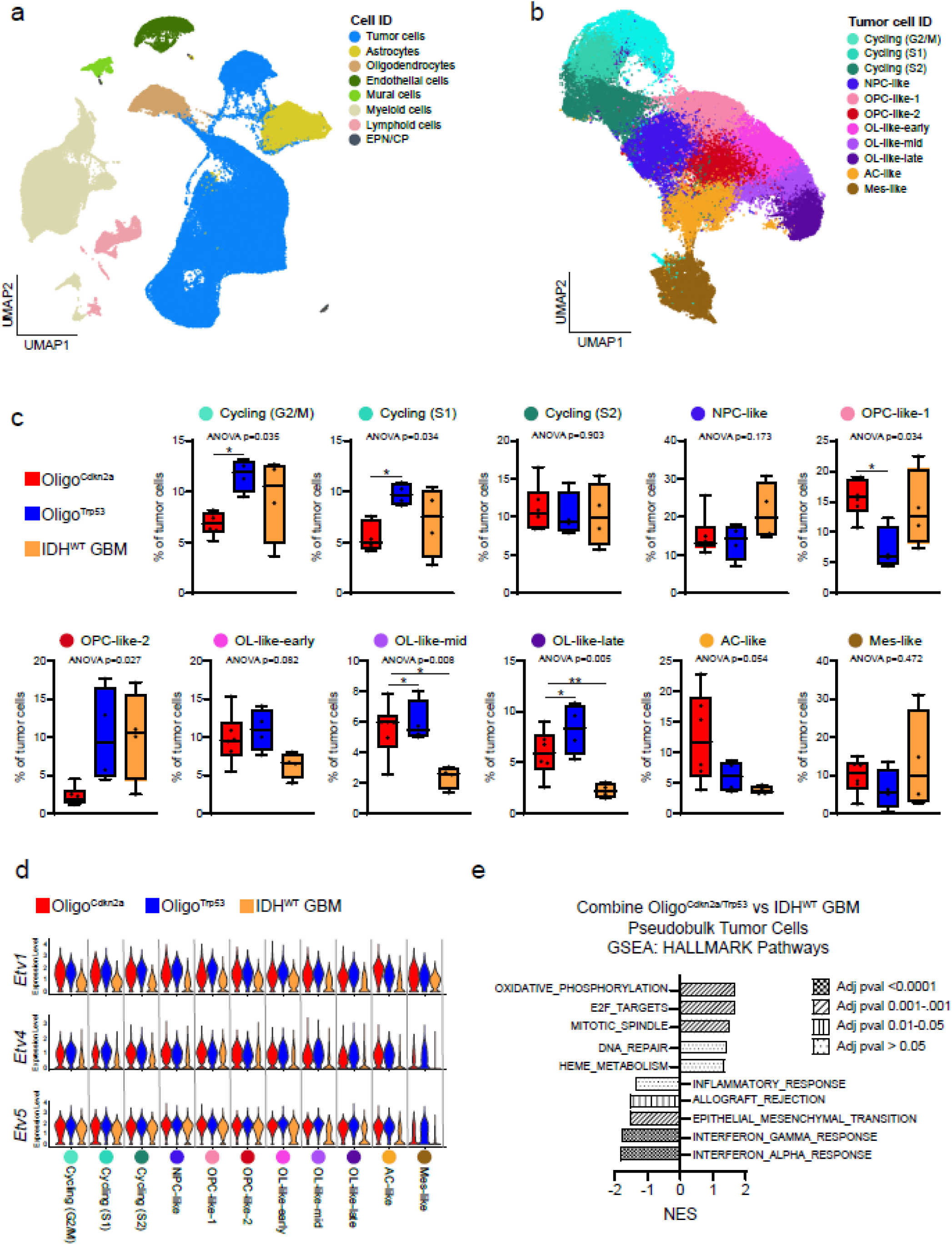
Single cell profiling identifies tumor cell-state and transcriptional heterogeneity driven by tumor genetics. **a,** UMAP visualization of cells in tumor samples colored by cell-type. **b,** UMAP visualization of tumor cells colored by cell-state programs. **c,** Frequency plots showing the contribution of tumor cell-state programs across conditions. Box plots show individual values, median (line), box limits (25th-75th percentiles), and whiskers (min-max). One-way ANOVA p-value displayed, and *p<0.05 or **p<0.01 denotes significance for Tukey’s multiple comparisons test. Oligo^Cdkn2a^ n=6, Oligo^Trp53^ n=4, IDH^WT^ GBM n=4. NPC: neural progenitor cell; OPC: oligodendrocyte progenitor cell; OL: oligodendroglioma; AC: astrocytic cell; Mes: mesenchymal. **d,** Violin plots showing *Etv1*, *Etv4* and *Etv5* expression in tumor cell-state clusters for each background. **e,** Gene set enrichment analysis (GSEA) of HALLMARK gene sets for pseudo bulk tumor cell comparison. The top five most significant (adj-pval) gene sets are plotted by normalized enrichment score (NES).

Pseudobulk analysis of tumor cell clusters was used to identify differentially expressed genes between conditions (logFC ≥ 0.7 or ≤ −0.7; adjusted p-value < 0.05). Known targets of CIC repression that are highly expressed in human IDH-O^29^ such as *ETV1*, *ETV4* and *ETV5* were consistently upregulated in Oligo^Cdkn2a^ and Oligo^Trp53^ tumor cell clusters when compared to IDH^WT^ GBM (**Figure 5d**). Next, gene set enrichment analysis (GSEA) was performed on pseudo bulk profiles of all tumor cell clusters to identify pathways enriched in specific conditions. When Oligo^Cdkn2a^ and Oligo^Trp53^ tumor cells were combined, they demonstrated enrichment of HALLMARK pathways for oxidative phosphorylation, E2F targets, and mitotic spindle, whereas IDH^WT^ GBM tumor cells showed increased interferon-α/γ response, epithelial mesenchymal transition (EMT) and inflammatory pathway activity (**Figure 5e**). Oligo^Cdkn2a^ tumor cells showed enrichment of HALLMARK pathways such as TNF-alpha signaling via NF-kappa beta, p53, and hypoxia, relative to both *Trp53*-mutant conditions combined, which showed increased MYC and interferon-α/γ response pathway activity (**Supplementary Figure 5e**). Oligo^Trp53^ tumor cells were specifically enriched in proliferative pathways (e.g., MYC targets, E2F targets, G2M checkpoint) and interferon alpha/gamma response compared to other conditions (**Supplementary Figure 5e**).

Thus, single-cell analyses revealed that Oligo^Cdkn2a^ and Oligo^Trp53^ tumors are enriched for oligodendrocyte-lineage cell states and CIC target gene expression, with *Trp53* loss further promoting highly proliferative, progenitor-like programs. At a pathway level, metabolic reprogramming is present in both Oligo^Cdkn2a^ and Oligo^Trp53^ tumors. Oligo^Cdkn2a^ tumor cells preferentially activated p53, hypoxia, and TNF-α/NF-κB pathways, whereas Oligo^Trp53^ and IDH1^WT^ GBM tumor cells showed stronger MYC, interferon-α/γ, EMT, and inflammatory signaling.

### Tumor Genotype Shapes Heterogeneity in the Glioma Immune Microenvironment

Leveraging our single cell dataset, we next examined differences in the TME, focusing on the *Ptprc*+ (Cd45) immune cell compartment (**Supplementary Figure 6a**). Myeloid populations such as tumor associated microglia (*Csf1r*+ / *P2ry12*+) and macrophages (*Csf1r*+ / *P2ry12*-) represented the majority of immune cells in tumors (**Figure 6a-c**). Tumor associated microglia could be divided into three subclusters defined by expression of *Cxcl9* (microglia *Cxcl9*-high) and *Spp1* (microglia *Spp1*-high and microglia *Spp1*-low) (**Figure 6a-c**). Oligo^Cdkn2a^ tumors displayed an increased proportion of *Spp1*-high microglia compared to other tumor conditions (**Figure 6c** and **Supplementary Figure 6b, c**). Cycling tumor associated microglia/macrophages (TAMs) were significantly increased in Oligo^Trp53^ tumors (**Figure 6c** and **Supplementary Figure 6b, c**), indicating that *Trp53* status influences not only tumor cell-intrinsic programs but also other cellular interactions in the TME. Lymphocyte populations, including T cells, NK cells, and B cells, although comprising only a small proportion of the immune cell compartment, were more frequently observed in Oligo^Trp53^ and IDH^WT^ GBM tumors (**Figure 6c** and **Supplementary Figure 6b, c**). *Cd3e*+ T cell populations consisted of *Cd8a*+ cytotoxic T cells (Tc), *Cd4*+ helper T cells (Th) and *Cd4*+/*Foxp3*+ regulatory T cells (T-regs). Importantly, T cell populations displayed high expression of genes associated with exhaustion/dysfunction, including *Pdcd1* (Pd1), *Ctla4* and *Tox* (**Figure 6d**), mirroring exhausted phenotypes found in human gliomas^43–45^.

**Figure 6.**
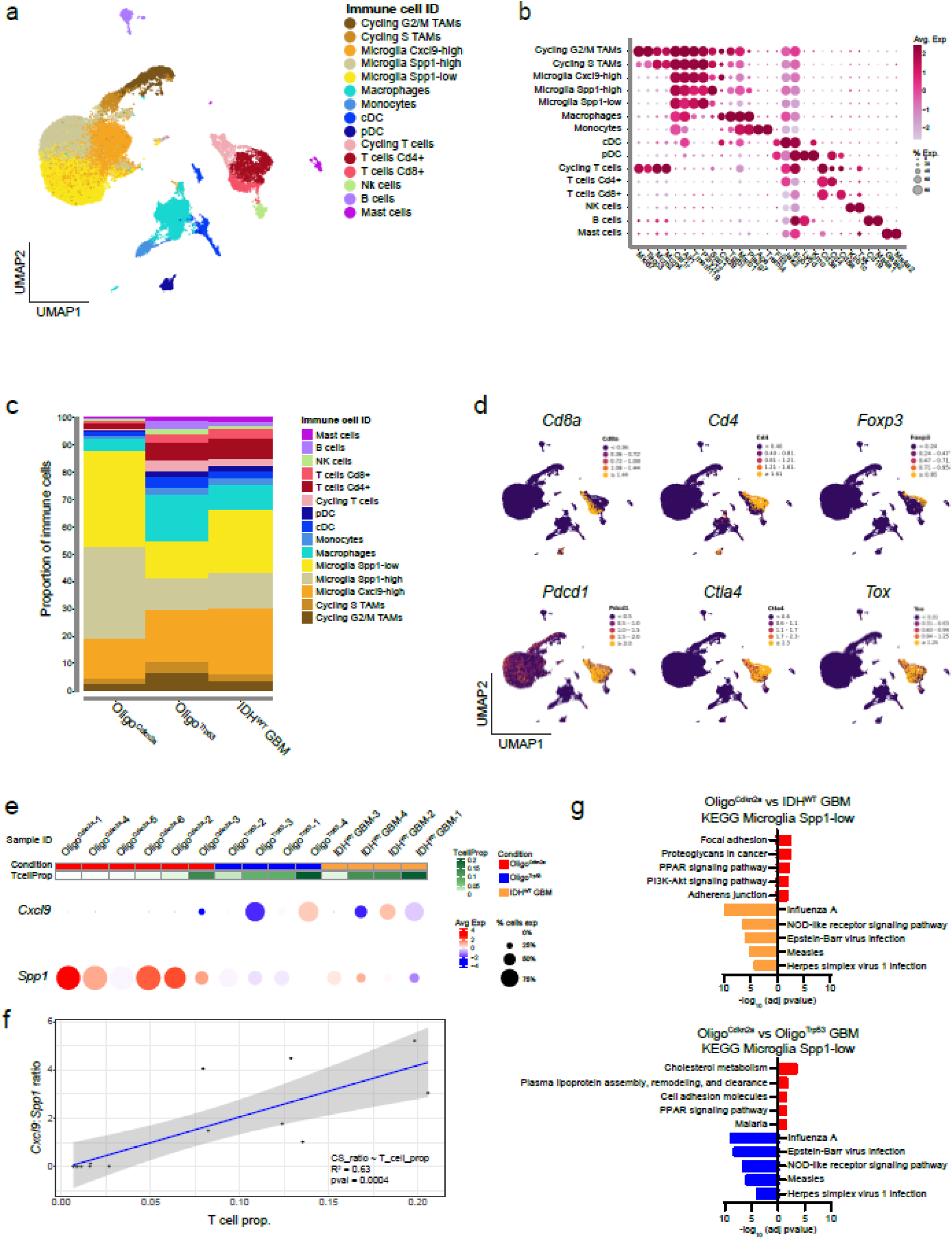
Tumor Genotype Shapes Heterogeneity in the Glioma Immune Microenvironment. **a,** UMAP visualization of *Ptprc*+ (i.e., Cd45+) immune cells colored by cell-state programs. **b,** Dot plot displaying the relative expression of cell-type specific markers in the designated cell-types. **c,** Stacked bar plot illustrating the relative proportion of immune cell populations across backgrounds**. d,** UMAP feature plots displaying the expression of T cell lineage markers (*Cd8a*, *Cd4*, *Foxp3*) and exhaustion-associated markers (*Pdcd1*, *Ctla4*, *Tox*). **e,** Dot plot displaying *Cxcl9* and *Spp1* expression in both microglia and macrophage populations across individual tumor samples. Samples are annotated by genotype and ranked by the proportion of T cells within the immune cell population as shown. **f,** Correlation between the *Cxcl9*:*Spp1* expression ratio and the proportion of infiltrating T cells across tumor samples. Statistical significance was determined by linear regression (R^2^ = 0.63, p = 0.0004). **g,** ORA of KEGG gene sets found in DEGs based on pseudo bulk comparison of immune cell-type clusters. The top five most significant gene sets are plotted by −log_10_(adj-pvalue).

Previous work on interpatient variation in macrophage polarity programs identified a *CXCL9*:*SPP1* axis regulated by T cell-derived IFN-γ and hypoxia^46^. Because each IUE derived tumor arises independently *in situ*, we asked whether similar intertumoral heterogeneity could be found across murine tumors. To investigate this, we first plotted the proportion of T cells against the expression of *Cxcl9* and *Spp1* in the tumor associated microglia and macrophage clusters. From this, a pattern across samples emerged, where increased T cell presence was linked to elevated *Cxcl9* and decreased *Spp1* expression (**Figure 6e**). We then plotted the ratio of *Cxcl9*:*Spp1* expression against the proportion of T cells and found a significant correlation (**Figure 6f**). Thus, dynamic interplay between immune cell components can influence overall TME profiles in murine glioma models, capturing not only variation between genetic conditions but also at the individual tumor level.

Finally, we performed pseudobulk analysis of immune cell clusters to identify differentially expressed genes (logFC ≥ 0.7 or ≤ −0.7; adjusted p-value < 0.05) and enriched pathways between conditions. Immune activation related pathways were enriched in microglia and macrophage clusters in *Trp53*-mutant conditions (Oligo^Trp53^ and IDH^WT^) when compared to Oligo^Cdkn2a^, further validating their activity in these more aggressive tumor conditions (**Figure 6g** and **Supplementary Figure 7a-c**). Oligo^Cdkn2a^ microglia and macrophages showed enrichment in cholesterol metabolism, PPAR signaling, and proteoglycan regulation (**Figure 6g** and **Supplementary Figure 7a-c**). Minimal differences between Oligo^Trp53^ and IDH^WT^ GBM immune clusters again verified their shared activation state (**Supplementary Figure 7a**). Together, these findings show that tumor genotype not only shape intrinsic tumor cell programs but also drive distinct, heterogeneous immune microenvironments, with *Trp53*-mutant tumors exhibiting heightened inflammatory states.

## Discussion

The absence of faithful mouse models has been a major obstacle to studying IDH-O. No existing IDH-mutant mouse glioma models incorporate *Cic* loss, despite its predicted importance in IDH-O. Here, we describe the generation of a murine IDH-O model that integrates IDH-mutation, loss of 1p/19q targets (*Fubp1* and *Cic* respectively) and other relevant genetic alterations to drive gliomagenesis. By testing different genetic combinations, we find that the Oligo^Cdkn2a^ mouse model recapitulates key features of IDH-O across multiple domains. Classic oligodendroglioma histopathology is preserved, including round uniform nuclei with hydropic cytoplasm. Tumors display metabolic alterations consistent with IDH-mutation, including D-2HG accumulation, and exhibit enhanced differentiation along the oligodendrocyte lineage trajectory. Expression of transcriptional targets and programs upregulated in IDH-O are similarly increased, reflecting strong transcriptional fidelity. Together, these features demonstrate that the Oligo^Cdkn2a^ mouse model authentically mirrors the phenotypic, metabolic, and molecular landscape of human IDH-O.

We found that Oligo^Cdkn2a^ and Oligo^Trp53^ mouse models displayed biases towards oligodendrocyte lineage differentiation, suggesting a genetic determinant for this cell-state program. Prior work has demonstrated the necessity of RAS/MAPK signaling in promoting glial lineage identity and gliomagenesis, and *Cic* loss in promoting neural stem cell proliferation and OPC cell-state programs^47,48^. Our data is in line with these prior findings and suggest that *Cic* loss not only supports the maintenance of glial stem/progenitor cells but also primes tumor cells to differentiate along the oligodendrocyte axis. We also show that *Trp53* loss, which is not a common occurrence in human IDH-O tumors, alters tumor pathogenesis, significantly decreasing latency, and modifying histological and transcriptional features. To the best of our knowledge, this is the first experimental evidence that angulated nuclei with dense clumped chromatin are a morphologic manifestation of *TP53* inactivation in gliomas. Oligo^Trp53^ tumors also showed a propensity for enhanced immune cell infiltration and inflammatory signatures, similar to that found in an IDH^WT^ GBM tumor model. These data would support prior findings suggesting that *TP53* mutation status contributes to TME differences between IDH-A (*TP53* mutant) and IDH-O (*TP53* wild-type)^29^. More broadly, comparison of distinct murine glioma subtype models affords an opportunity to examine features of the human disease. For example, supratentorial human GBM is known to be highly invasive, with tumor cells reaching hindbrain structures at advanced stages of disease^49^. These unique genetic mouse models recapitulated aspects of this behavior, as IDH^WT^ GBM tumors showed highly invasive characteristics, including the presence of GFP+ tumor cells in the brainstem, whereas this type of invasion was not present in Oligo^Cdkn2a^ and Oligo^Trp53^ models. This also aligns with our patient-based data, wherein GBMs require much less time than IDH-mutant gliomas to reach similar levels of disease burden in the brainstem^44^.

The Oligo^Cdkn2a^ and other IUE brain tumor models presented here provide a valuable platform to investigate mechanisms of tumor development, interactions with the microenvironment, and test new therapeutic strategies. The modular nature of IUE offers a flexible system to assess different genetic combinations. By integrating both gain- and loss-of-function approaches, the IUE platform offers a versatile framework to examine how specific genetic alterations influence tumor development and therapy response. For example, investigating *NOTCH1* mutations in Oligo^Cdkn2a^ mouse models could provide additional insight into how these mutations alter tumor cell-state programs and response to IDH inhibition^50^. Additional studies on the function of *FUBP1*, recently identified as general splicing factor that facilitates 3’ splice site recognition^51^, are also warranted to delineate its role in gliomagenesis. Importantly, Oligo^Cdkn2a^, Oligo^Trp53^ and IDH^WT^ glioma mouse models are immune competent and display diverse tumor immune microenvironment interactions that mirror features of human tumors. We anticipate that these models will serve as valuable tools to investigate TME interactions and for testing new targeted and immunotherapy approaches, similar to studies in other brain tumor types that leverage IUE and IUE-derived models^52–57^.

## Methods

### In utero electroporation

All mouse experiments were conducted in accordance with the Institutional Animal Care and Use Committee protocols (IACUC # 22-05-15-01 and 25-05-05-01) at the University of Cincinnati. Time-pregnant CD-1 IGS (Charles River, #022) were used for all experiments. In utero electroporation (IUE) was performed as previously described^58^. Approximately, 1 µL of a DNA plasmid / mRNA mixture containing 0.05% Fast Green was injected into the lateral ventricle of e13.5– 14.5 embryos using a pulled glass capillary pipette. The final DNA/RNA solution consisted of DNA plasmids (each plasmid at 1µg/µL), 0.33mM Cas9 (TriLink; CleanCap Cas9 mRNA # L-7206-100) and 20µM sgRNA (Editco; mouse gene knockout kit). Injected embryos were electroporated by applying 5 square pulses (45V, 50-ms pulses with 950 ms intervals) with 3-mm tweezer electrodes directed toward the dorsal cortex (ECM830 BTX/Harvard Bioscience). Embryos were returned to the abdominal cavity, incision was sutured, and the female was monitored until fully recovered. Subsequently, successfully electroporated offspring were monitored regularly, and mice presenting symptoms related to tumor burden were euthanized according to protocols approved by the Institutional Animal Care and Use Committee at the University of Cincinnati.

### Plasmid Construction and sgRNAs

The Piggybac donor plasmid PBCAG-Ires-eGFP was used to engineer IDH1^R132H^ and PIK3CA^E545K^ plasmids. Gene cDNA sequences for IDH1^R132H^ (Addgene, #81686) and PIK3CA^E545K^ (Addgene, #82881) were PCR-amplified from template plasmids and inserted into EcoRI-linearized pBCAG-Ires-eGFP backbone using the In-Fusion Snap Assembly cloning kit (Takara Bio, #639298). The resulting constructs were transformed and propagated in Stellar competent bacteria cells (Takara Bio, #636766). Plasmid DNA was subsequently isolated using an endotoxin-free purification protocol (Macherey-Nagel, #NC1688968). Plasmid sequences were verified by whole-plasmid sequencing (Plasmidsaurus). DNA plasmids used include: PBCAG-IDH1^R132H^-Ires-eGFP, PBCAG-PIK3CA^E545K^-Ires-eGFP, PBCAG-eGFP, pCAG-PBase.

Mouse gene knockout kits (Editco) included the following sgRNA sequences for each gene target: *Cic:* 5′-AAAUGUCGAACCUCGUUCUG-3′, 5′-GCUGCACAGAGGGGUCGGGC-3′, and 5′-UGCUCUGGUUGGAGGCCACA-3′; *Fubp1*, 5′-AGACGACGGCGGAGGCACUG-3′, 5′-UACCUGCCGCGCUCUCUGCA-3′, and 5′-CGGCUCUCGGGGGUCGGGGA-3′; for *Cdkn2a*, 5′-GCAGCACCACCAGCGUGUCC-3′, 5′-CGGUGCAGAUUCGAACUGCG-3′, and 5′-GACUUUUCAGGUGAUGAUGA-3′; for *Trp53*, 5′-AGACGUGCCCUGUGCAGUUG-3′, 5′-GAAGUCACAGCACAUGACGG-3′, and 5′-UGAGGGCUUACCAUCACCAU-3′; for *Nf1*, 5′-ACCAACAUGCAGCCGAACUU-3′, 5′-UCUUUAGUCGCAUUUCUACA-3′, and 5′-AAUUCUUGUCUGCAUAAGAG-3′; and for *Pten*, 5′-AAUCCCAUAGCAAUAAUAUU-3′, 5′-ACAAUAUUGAUGAUGUAGUA-3′, and 5′-CAAAUACUGGUCUGUGUUUU-3′.

### Genomic sequencing of CRISPR-Cas9 editing

Sequencing for CRISPR-Cas9 mediated insertions and deletions (INDELs) was performed on genomic DNA (gDNA) extracted from GFP+ tumor tissue and cell lines generated from tumor tissue (Zymo Research, #D7003). DNA primers (IDT) surrounding the CRISPR-Cas9 cut sites were used to PCR amplify genomic locus. DNA primer pairs for each gene are as follows: *Cic*, 5′-CCATGGTATGGGTTCTGAAGC-3′ and 5′-CTCTCAGGGCACACTGCTC-3′; *Fubp1*, 5′-ATTTCCGTCTGCCAGTCTCC-3′ and 5′-CTGAGCGCCCAGAACCTTC-3′; *Cdkn2a*, 5′-GAAGTATAACATTCCAGAAAGACTAGG-3′ and 5′-TTCCCAGCGGTACACAAA-3′; *Trp53*, 5′-TGGTGCTTGGACAATGTGTT-3′ and 5′-GCTGTGGCGAAAAGTCTGC-3′; *Nf1*, 5′-TGTTTGCAGGTAAAGGAAAAGCC-3′ and 5′-AGAGAGAGATGGCCATGGAGA-3′; *Pten*, 5′-TGTCCAGGCTATAGCTCAAGG-3′ and 5′-CCAGAGAATAACCTGGTAAGAGACA-3′. PCR amplicons were gel-purified using a NucleoSpin Gel and PCR Clean-up kit (Macherey-Nagel, #NC0389462) and submitted for Sanger sequencing at the Cincinnati Children’s Hospital Medical Center (CCHMC) DNA Core using the following sequencing primers: *Cic:* 5′-CATGGTATGGGTTCTGAAGCTTG-3′; *Fubp1:* 5′-GATGGTCGTGCAAGAATGTGATAGAG-3′; *Cdkn2a:* 5′-TTCCCAGCGGTACACAAA-3′; *Trp53:* 5′-GTGCTTGGACAATGTGTTTCATTAGT-3′; *Nf1:* 5′-ATGTGATTATTTCCATTTTAGCAACCAAAG-3’; *Pten:* 5′-CAGACAAGTACATTTGACGACCTTTTATA-3′. Sanger sequencing trace files from control and experimental conditions were analyzed using the Inference of CRISPR Edits (ICE) portal (Editco).

### Tissue collection, histological processing, and immunostaining

Brains were collected as previously described^58,59^. For histopathology brain samples were fixed overnight in 10% formalin (Azer Scientific, #PFNBF-20) and then transferred into 70% ethanol before being processed for paraffin embedding. Five-micrometer-thick sections were prepared on a microtome and processed for hematoxylin-eosin staining. Stained slides were scanned at 40 on an Aperio digital slide scanner. Immunohistochemistry (IHC) for noted antigens was performed at the CCHMC pathology core using the automated Ventana BenchMark stainers. Antibodies included: IDH1 R132H (CellMarque, #456R-38) and GFP (Invitrogen, #A11122). For immunofluorescent stains, brains were rapidly dissected in ice-cold phosphate buffered saline (PBS) and then fixed in 4% paraformaldehyde overnight. Fixed brains were washed in cold PBS for 1 hour twice before transferring to a 30% sucrose/PBS solution at 4°C overnight. Once sunk brains were embedded in tissue freezing media. Free-floating sections 50 μm thick were made using a cryostat (Leica). Free-floating sections were transferred to blocking solution (PBS + 0.5% Triton X-100 + 10% normal donkey serum) at room temperature for 30–60 minutes before adding specific combinations of antibodies. Primary antibodies were incubated at 4°C overnight. The next day, sections were washed in PBS and then transferred to blocking solution containing the appropriate secondary antibodies (1:1,000) and incubated at 4°C overnight. Finally, sections were washed in PBS followed by a 10-minute incubation in Hoechst (1:1000 in PBS) and washed in PBS again before being mounted onto slides (Fisher, Superfrost) and coverslipped (Prolong Gold Antifade, ThermoFisher). Primary antibodies used included: eGFP (Aves, #GFP1020), Olig2 (Millipore, #Ab9610), Gfap (Cell Signaling, #12389), Ki67 (Cell Signaling, #9129), Corresponding secondary antibodies used were all purchased from Invitrogen. Images were acquired on a confocal microscope (Nikon A1), and image analysis was performed in FIJI / Image J (National Institutes of Health).

### Tumor dissociation

Brain were collected in ice-cold PBS and GFP-positive tumor regions were micro-dissected under a fluorescent stereoscope. GFP-positive tissues was minced with scalpels before enzymatic digestion in a papain solution. Papain (Worthington, #LS003126) solution was prepared by dissolved the enzyme (20 units/mL) in NeuroCult basal media, followed by addition of N-acetyl-L-cysteine (Sigma, #A9165-25G) and DNaseI (Worthington, #LS002007). Samples were incubated for 30 min at 37°C and then mechanically triturated to dissociate before passing through a 30µm cell strainer. Myelin and cellular debris were removed by density gradient centrifugation using Debris Removal Solution (Miltenyi, #130-109-398) according to the manufacturer’s protocol. Erythrocytes were subsequently eliminated using ACK Lysing Buffer (Gibco, #A10492-01).

### Primary cell cultures

Primary murine glioma cells derived from dissociated Oligo^Cdkn2a^, Oligo^Trp53^, and IDH^WT^ GBM tumors were grown in NeuroCult basal medium containing 10% NeuroCult Proliferation Supplement (Stemcell Technologies, #05702) and supplemented with 2µg/mL heparin (Stemcell Technologies #07980) plus growth factors (Shenandoah Biotech #PB-500-20) including 20ng/mL Epidermal Growth Factor (EGF; #200-53), 20ng/mL Fibroblast Growth Factor-basic (FGF; #200-12), 10ng/mL PDGF-AA (#200-54) and 10ng/mL PDGF-BB (#200-58). Cultures were grown as adherent monolayers on tissue culture treated flasks coated with laminin (Sigma #L2020). For metabolomic studies cells were grown in Neurobasal A medium supplemented with B27, N2 and the above described growth factors.

### Nucleic acid isolation

Total RNA and gDNA were co-extracted from micro-dissected GFP+ primary tumor tissue or murine glioma cell lines using the Quick-DNA/RNA Miniprep Plus Kit (Zymo Research, #D7003) according to the manufacturer’s protocol. To preserve the nucleic acid integrity, freshly harvested samples were immediately stabilized in DNA/RNA Shield (Zymo Research, #D7003) prior to processing.

### Metabolic Profiling

Cells were treated with vehicle (DMSO) or vorasidenib (1 uM) for 72 h. Metabolites were extracted by dual phase extraction using methanol-chloroform-water as described earlier^39^. 1H-MRS spectra (1D water presaturation ZGPR sequence, 90 FA, 3s TR, 256 acquisitions) were acquired using a 600 MHz Avance spectrometer (Bruker BioSpin). Metabolites were quantified by normalizing to a trimethylsilyl propanoic acid reference of known concentration and to cell number and correcting for saturation and Nuclear Overhauser effect.

### Patient genetic dataset

Whole-genome sequencing data from 62 IDH-mutant 1p/19q codeleted oligodendroglioma patients were accessed as part of the OligoNation / CBTN provisional dataset deposited in pedscbioportal (https://pedcbioportal.kidsfirstdrc.org/study/summary?id=684c727fb62c3850c29fe389).

### Bulk RNA-sequencing

The directional polyA RNA-seq was performed by the Genomics, Epigenomics and Sequencing Core at the University of Cincinnati using established protocols as previously described^60,61^ with updates. Total RNA was isolated as noted above using the Quick-DNA/RNA Miniprep Plus Kit (Zymo Research, #D7003). The quality of total RNA was determined by Bioanalyzer (Agilent, Santa Clara, CA). Briefly, 500 ng of high-quality total RNA per sample was subjected to poly(A) enrichment using the NEBNext Poly(A) mRNA Magnetic Isolation Module (New England Biolabs). Sequencing libraries were subsequently constructed using the NEBNext Ultra II Directional RNA Library Prep Kit for Illumina (New England Biolabs) with unique dual indexing, undergoing amplification for 9 PCR cycles. Final libraries were evaluated for quality, quantified via Qubit fluorometry (Thermo Fisher Scientific), and normalized. Paired-end sequencing (2 × 61 bp) was performed on a NextSeq 2000 platform (Illumina), achieving a sequencing depth of approximately 30 million read pairs per sample. Sequence reads were aligned to the reference genome using the STAR aligner, and reads aligning to each known transcript were counted using Bioconductor packages for next-generation sequencing data analysis. The differential expression analysis between different sample types was performed using the negative binomial statistical model of read counts as implemented in the edgeR Bioconductor package. Transcriptional profiles were interrogated with both Interactive Gene Expression Analysis Kit (for microarray and RNA-seq data), an R (v3.3.2), and JavaScript-based open-source desktop application^62^ and integrated Differential Expression and Pathway analysis^63^ tools. Functional enrichment of differentially expressed gene lists between tumor groups was performed with gProfiler^64^ using default settings. All RNA-seq files are deposited in Gene Expression Omnibus as GSE326763.

### GEM-X Flex sample preparation and sequencing

Mouse tumor samples were collected as described in the tissue dissociation section. Following dissociation, single-cell suspensions were resuspended in single-cell buffer (DPBS + 0.04% BSA) to quantify cell density and verify viability >90% using the automated LUNA-FL Dual Fluorescence Cell Counter (Logos Bio, Annandale, VA). Approximately 1 million cells were fixed using the GEM-X Flex Sample Preparation v2 kit (10x Genomics, #PN-1000781) according to the manufacturer’s protocol. Samples were fixed overnight and then transferred into quenching buffer containing 10% glycerol for long-term storage at −80°C. Prior to library preparation, samples were thawed and adjusted to the recommended 300,000 cells per sample. Following hybridization with the GEM-X Flex probe set and the pooled wash workflow, approximately 14,500 cells were loaded per lane for GEM generation, targeting recovery of 10,000 cells per sample.

After GEM generation, ligation, preamplification, and library indexing were performed according to the GEM-X Flex protocol. Library quality and fragment size distribution were assessed using a Bioanalyzer (Agilent, Santa Clara, CA). High-quality libraries were sequenced on a NovaSeq X system (Illumina, San Diego, CA) using a read 28|10|10|90 configuration (Read 1 | Index 1 | Index 2 | Read 2), generating approximately 40,000 read clusters per cell for a targeted recovery of 10,000 cells per sample.

### Preprocessing, alignment, and cell calling

Sequencing data were processed using Cell Ranger (v9.0.0, 10x Genomics). Reads were aligned to the *Mus musculus* reference genome mm10 with the corresponding 10x Genomics GEM-X Flex mouse probe set, and gene-by-cell UMI count matrices were generated for downstream analysis. The Cell Ranger cell-calling algorithm employs the order of magnitude (OrdMag) approach to estimate the number of recovered cells, calling barcodes as cells if total UMI counts exceed m/10, where m is the 99^th^ percentile of total UMI counts among the top N barcodes. To identify cells with low RNA content that may not be called by OrdMag, Cell Ranger additionally applies the EmptyDrops algorithm^65^, which detects significant deviations from the ambient RNA background model.

### Single-cell quality control, integration, and clustering

Cell Ranger outputs were analyzed using Scanpy. Quality control filtering included exclusion of cells with >5% mitochondrial transcript content. The relationship between the number of detected genes and the number of UMIs per cell was modeled using linear regression, and outliers were removed based on this relationship. Putative doublets were identified and removed using the scDblFinder algorithm with a probability threshold >0.3. Data integration across samples was performed using Harmony with numGenes = 2000 and normalization = log Normalized. PCA was performed, and 30 principal components were used for downstream dimensionality reduction and clustering. UMAP was computed with a minimum distance of 0.3 using a cosine metric. Clusters were identified using the Leiden algorithm with a resolution parameter of 0.8. Cell types were annotated by canonical markers.

### Pseudo bulk differential expression and functional analysis

Pseudo bulk differential expression analysis between tumor groups was performed using the pseudo bulk limma-voom workflow. Functional enrichment of differentially expressed gene lists was performed using gProfiler^64^ with default parameters. Ranked gene set enrichment analysis (GSEA) was performed on the differential expression results using the fgsea R package and the mouse Hallmark gene set collection from the Molecular Signatures Database^66^.

Single cell transcriptional profiling data files are deposited in Gene Expression Omnibus as GSE326990 and GSE326991.

### Statistics

Statistical analyses were performed in GraphPad Prism (v11.0.0). Statistical significance was determined by log-rank Mantel–Cox test, one-way ANOVA with Tukey’s multiple comparisons test, or Two-way ANOVA with Sidak’s multiple comparisons test, as described in the figure legends. P values < 0.05 were considered significant.

## Acknowledgement

We are grateful to the University of Cincinnati Laboratory Animal Medical Services (LAMS) for their assistance with animal housing, husbandry, and veterinary care. Bulk and single cell RNA-sequencing was performed by the Genomics, Epigenomics and Sequencing Core (GES Core), University of Cincinnati College of Medicine. Sanger sequencing was performed by the Genomics Sequencing Facility / Gene Analysis Core at Cincinnati Children’s Hospital Medical Center, and this project was supported in part by NIH P30 DK078392 (Gene Analysis Core of the Digestive Diseases Research Core Center in Cincinnati). The Cincinnati Children’s Integrated Pathology Research Facility (RRID: SCR_022637), was used for paraffin processing and embedding, as well as hematoxylin and eosin (H&E) and immunohistochemical (IHC) staining. This work was funded by the Oligo Nation Foundation, the UC Brain Tumor Center and the Holly and David Cook Family Brain Tumor Research Endowment (T.N.P) and supported by NIH grants CA120813, NS120547, and CA221747 (A.B.H).

## Ethics declarations

All mouse experiments were conducted in accordance with the Institutional Animal Care and Use Committee protocols (IACUC # 22-05-15-01 and 25-05-05-01) at the University of Cincinnati.

---

**Supplementary Figure 1.**
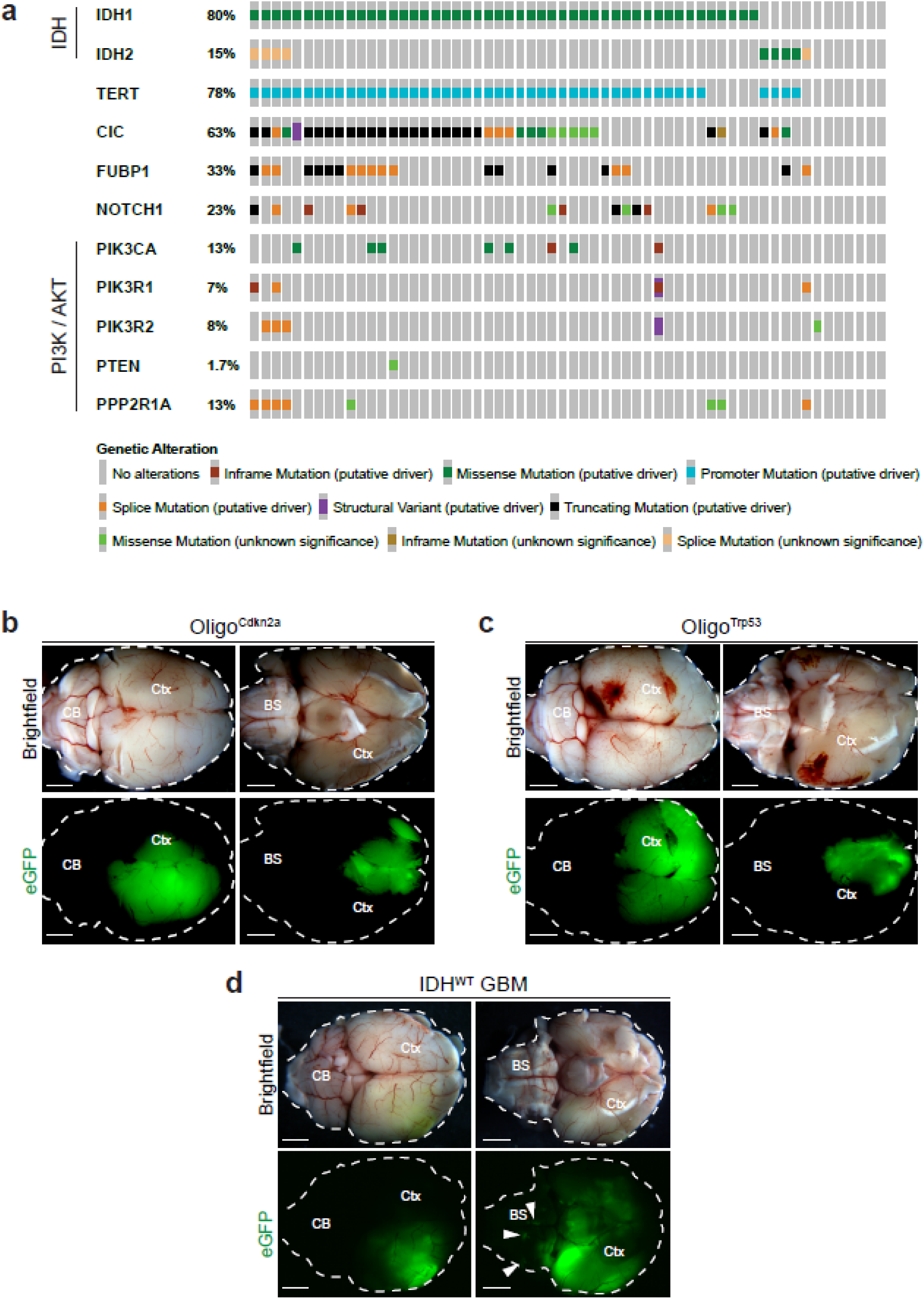
The genetic landscape of human IDH-mutant 1p/19q codeleted oligodendroglioma. **a,** Oncoprint displaying the top recurrent genetic alterations across 62 confirmed IDH-mutant 1p/19q oligodendroglioma patients (Oligo Nation/CBTN provision dataset). **b-d,** Brightfield and GFP whole brain images demonstrating the location of GFP-positive tumor cells across IUE conditions. Arrowheads denote spread of GFP-positive cells into the brainstem of IDH^WT^ GBM. Scale bar: 200µm. Ctx: cortex; CB: cerebellum; BS: brainstem.

**Supplementary Figure 2.**
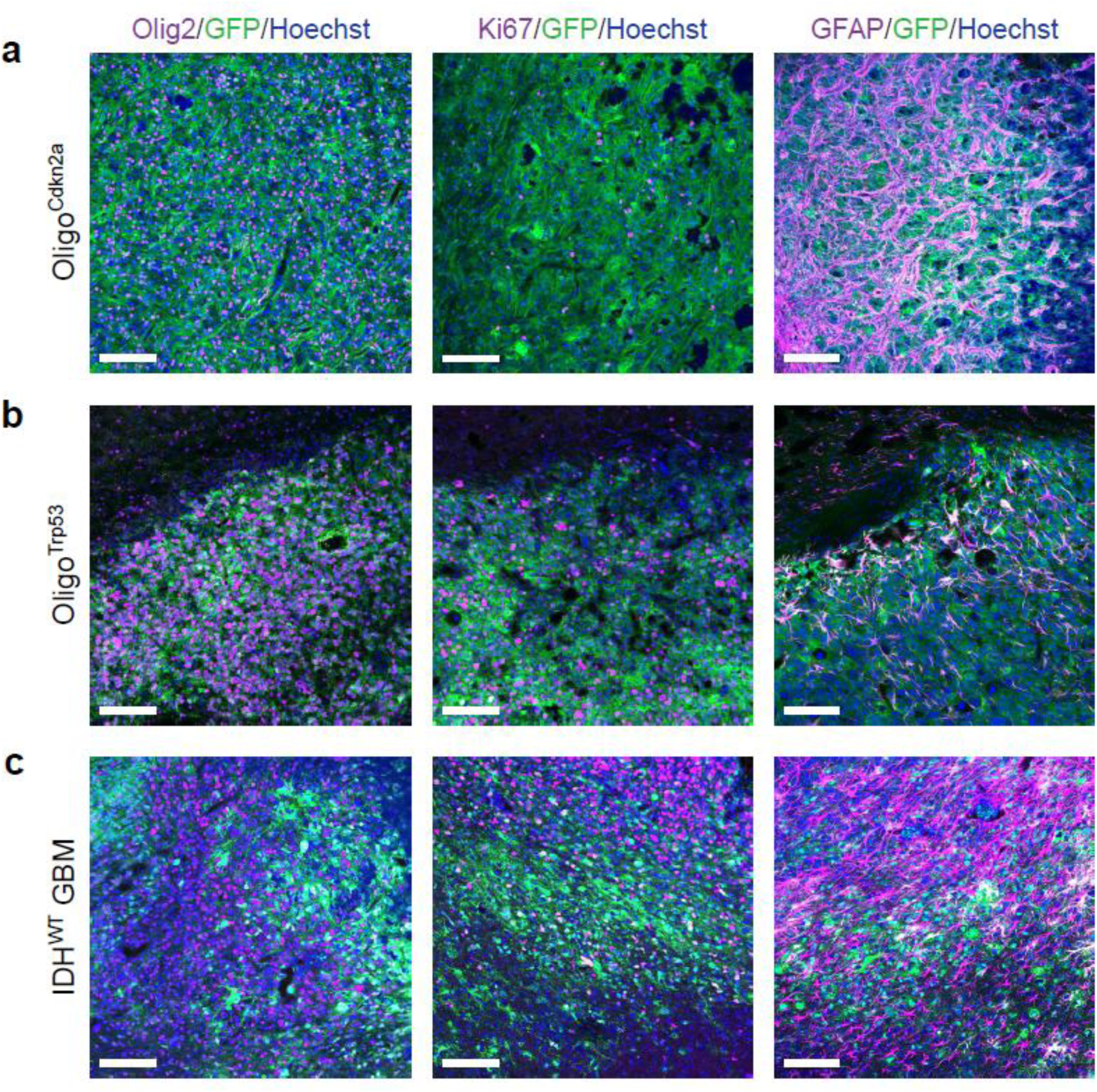
Shared expression of glioma markers in Oligo^Cdkn2a/^, Oligo^Trp53^ and IDH^WT^ GBM mouse models. **a,** Oligo^Cdkn2a^, **b,** Oligo^Trp53^, and **c,** IDH^WT^ GBM tumor samples co-stained for GFP (green), Hoechst (blue) and either Olig2, Ki67 or GFAP (magenta). Scale bar: 100µm.

**Supplementary Figure 3.**
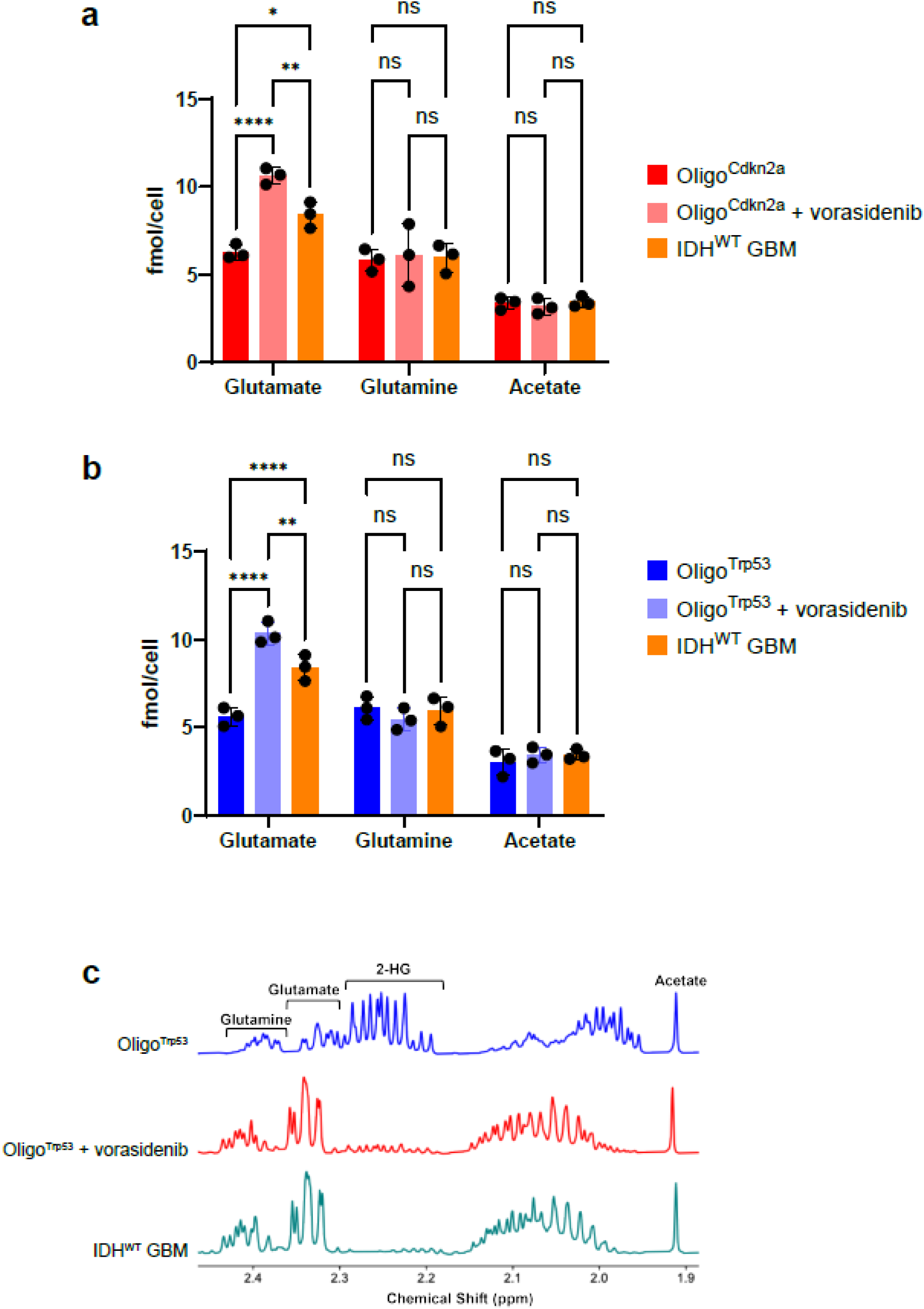
Metabolic profiling of oligodendroglioma models and response to IDH inhibition. **a,** Metabolite levels (fmol/cell) in Oligo^Cdkn2a^ cells treated with DMSO or vorasidenib (uM, 72 hrs) and IDH^WT^ GBM cells. **b,** Metabolite levels (fmol/cell) in Oligo^Trp53^ cells treated with DMSO or vorasidenib (uM, 72 hrs) and IDH^WT^ GBM cells. Values indicate mean ± SD; Two-way ANOVA with Sidak’s multiple comparisons test. *p<0.05, **p<0.005, ***p<0.001, ****p<0.0001. Oligo^Cdkn2a^ n=3, Oligo^Trp53^ n=3, IDH^WT^ GBM n=3. **c,** Representative ^1^H magnetic resonance spectra illustrating the metabolic profiles of Oligo^Trp53^ (top trace), vorasidenib-treated Oligo^Trp53^ (middle trace), and IDH^WT^ GBM (bottom trace) cell extracts. Peaks corresponding to glutamine, glutamate, 2-hydroxyglutarate (2-HG), and acetate are indicated across the 1.9–2.4 ppm chemical shift range.

**Supplementary Figure 4.**
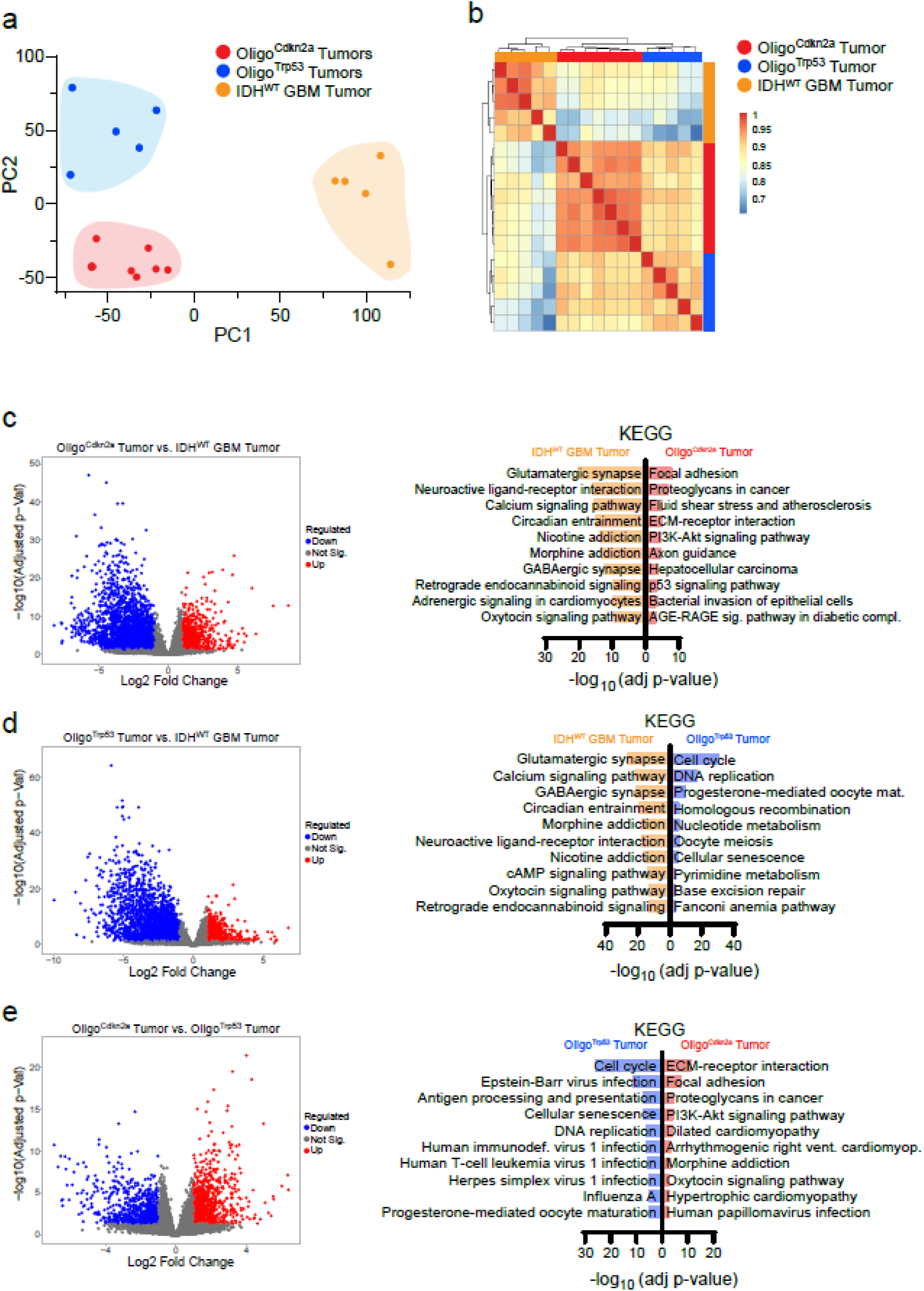
IDH- and Trp53-mutation status impact the transcriptional profile of tumor models. **a,** PCA plot of all tumor samples based on transcriptome-wide expression. **b,** Pearson’s correlation plot of all tumor samples. **c,** Volcano plot displaying significantly up- and down-regulated genes between Oligo^Cdkn2a^ vs IDH^WT^ GBM tumors, and ORA of KEGG gene sets in each condition. The top five most significant gene sets are plotted by −log_10_(adj-pvalue). **d,** Volcano plot displaying significantly up- and down-regulated genes between Oligo^Trp53^ vs IDH^WT^ GBM tumors, and ORA of KEGG gene sets in each condition. The top five most significant gene sets are plotted by −log_10_(adj-pvalue). **e,** Volcano plot displaying significantly up-and down-regulated genes between Oligo^Cdkn2a^ vs Oligo^Trp53^ tumors, and ORA of KEGG gene sets in each condition. The top five most significant gene sets are plotted by −log_10_(adj-pvalue).

**Supplementary Figure 5.**
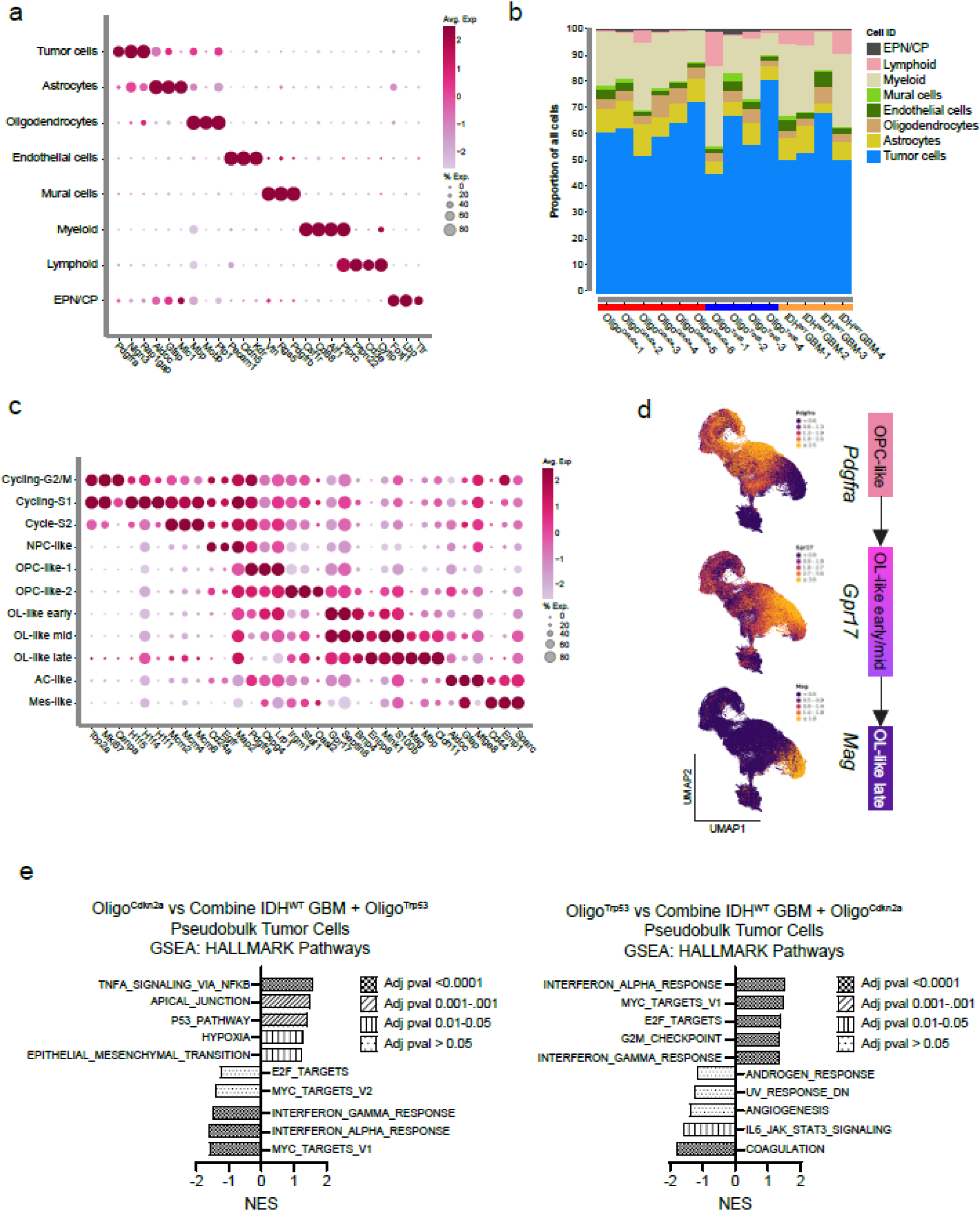
Single cell profiling identifies tumor cell-state and transcriptional heterogeneity driven by tumor genetics. **a,** Dot plot displaying the relative expression of cell-type specific markers in the designated cell-types. **b,** Stacked bar plot illustrating the relative proportion of immune cell populations in individual tumor samples. **c,** Dot plot displaying the relative expression of cell-type specific markers in the designated tumor cell-types. NPC: neural progenitor cell; OPC: oligodendrocyte progenitor cell; OL: oligodendroglioma; AC: astrocytic cell; Mes: mesenchymal**. d,** UMAP visualization of tumor cell clusters showing expression of selected oligodendrocyte lineage markers. **e,** GSEA of HALLMARK gene sets for pseudo bulk tumor cell comparison. The top five most significant (adj-pval) gene sets are plotted by normalized enrichment score (NES).

**Supplementary Figure 6.**
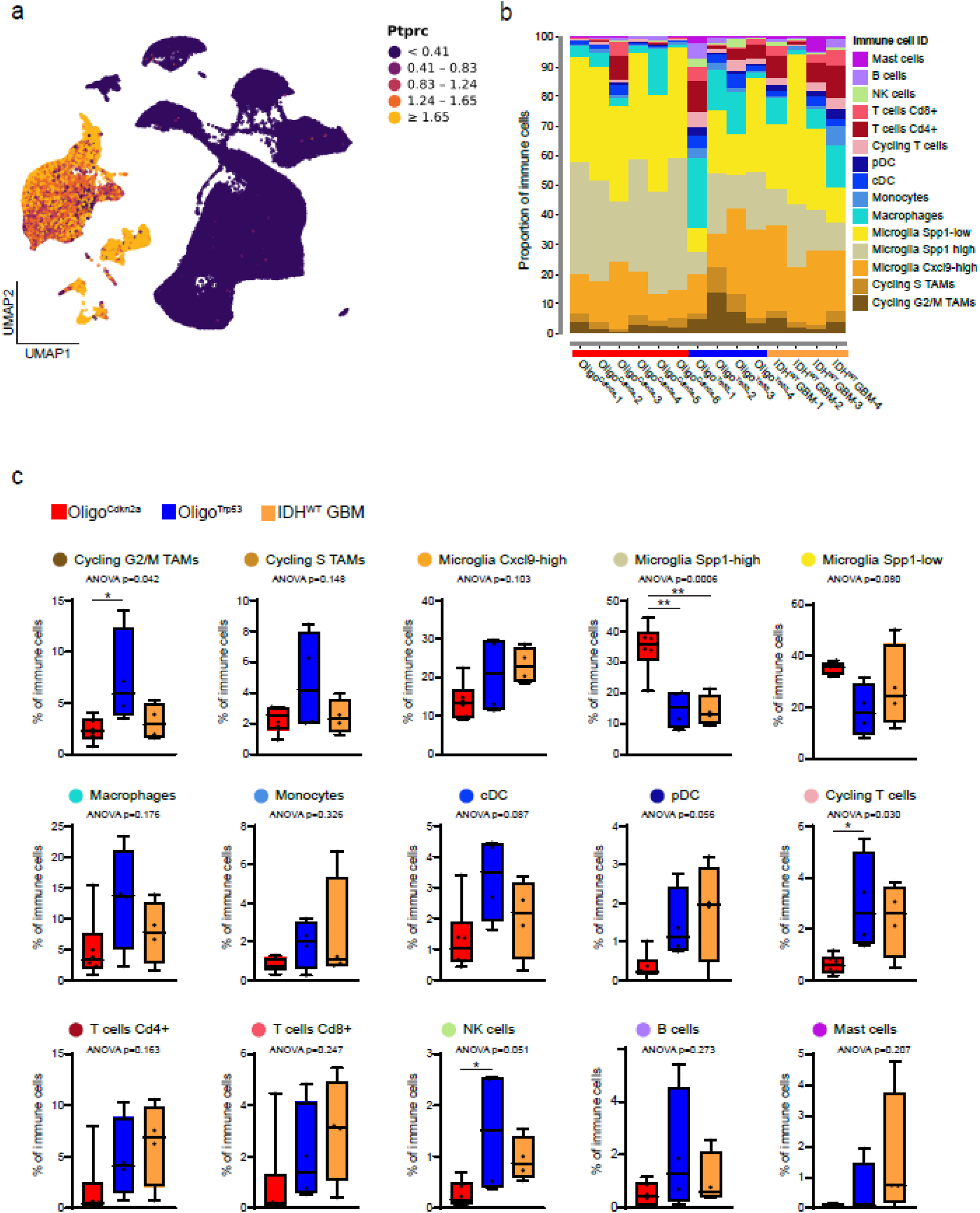
Tumor genotype shapes heterogeneity in the glioma immune microenvironment. **a,** UMAP visualization of all cells showing expression of *Ptprc* (Cd45+). **b,** Stacked bar plot illustrating the relative proportion of all major cell types in individual tumor samples. **c,** Frequency plots showing the contribution of immune cell-state programs across conditions. Box plots show individual values, median (line), box limits (25th-75th percentiles), and whiskers (min-max). One-way ANOVA p-value displayed, and *p<0.05 or **p<0.01 denotes significance for Tukey’s multiple comparisons test. Oligo^Cdkn2a^ n=6, Oligo^Trp53^ n=4, IDH^WT^ GBM n=4. TAMs: tumor associated microglia/macrophages; cDC: conventional dendritic cells; pDC: plasmacytoid dendritic cell; NK: natural killer cell.

**Supplementary Figure 7.**
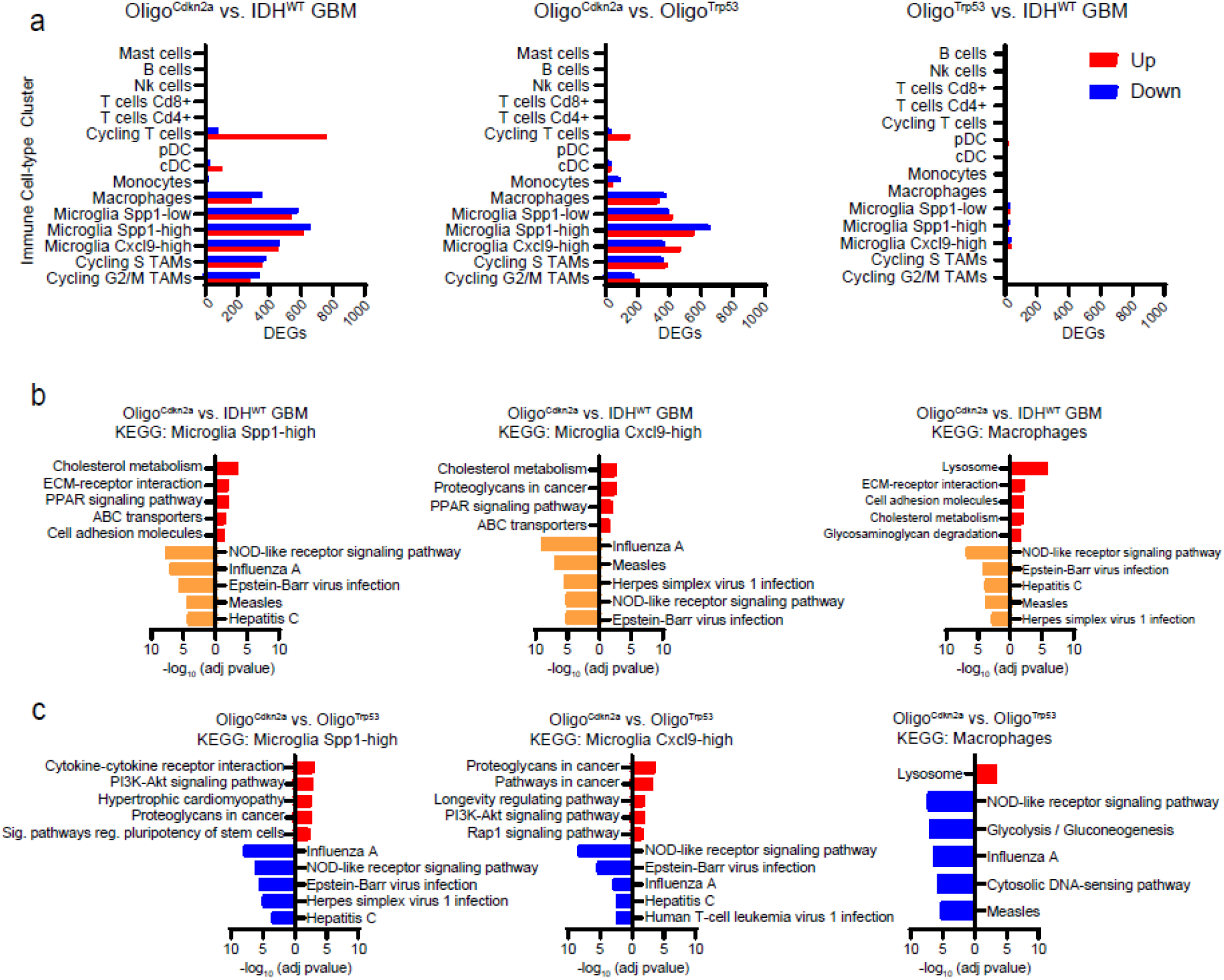
Tumor genotype shapes transcriptional heterogeneity in the glioma immune microenvironment. **a,** Bar plots showing number of differentially expressed genes (LogFC ≥ 0.7 or ≤ −0.7 and adj p-val < 0.05) identified in pseudo bulk comparisons for each immune cell-type cluster. **b,** Oligo^Cdkn2a^ vs IDH^WT^ GBM comparison, ORA of KEGG gene sets identified based on DEGs in noted immune cell-type cluster. Up to the top five most significant gene sets are plotted by −log_10_(adj-pvalue). **c,** Oligo^Cdkn2a^ vs Oligo^Trp53^ comparison, ORA of KEGG gene sets found in DEGs based on pseudo bulk comparison in noted immune cell-type cluster. Up to the top five most significant gene sets are plotted by −log_10_(adj-pvalue).

